# Global DNA methylation remodeling during direct reprogramming of fibroblasts to neurons

**DOI:** 10.1101/371427

**Authors:** Chongyuan Luo, Qian Yi Lee, Orly L. Wapinski, Rosa Castanon, Joseph R. Nery, Sean M. Cullen, Margaret A. Goodell, Howard Y. Chang, Marius Wernig, Joseph R. Ecker

## Abstract

Direct reprogramming of fibroblasts to neurons induces widespread cellular and transcriptional reconfiguration. In this study, we characterized global epigenomic changes during direct reprogramming using whole-genome base-resolution DNA methylome (mC) sequencing. We found that the pioneer transcription factor Ascl1 alone is sufficient for inducing the uniquely neuronal feature of non-CG methylation (mCH), but co-expression of Brn2 and Mytl1 was required to establish a global mCH pattern reminiscent of mature cortical neurons. Ascl1 alone induced strong promoter CG methylation (mCG) of fibroblast specific genes, while BAM overexpression additionally targets a competing myogenic program and directs a more faithful conversion to neuronal cells. Ascl1 induces local demethylation at its binding sites. Surprisingly, co-expression with Brn2 and Mytl1 inhibited the ability of Ascl1 to induce demethylation, suggesting a contextual regulation of transcription factor - epigenome interaction. Finally, we found that *de novo* methylation by DNMT3A is required for efficient neuronal reprogramming.

---

Mesoderm originated fibroblast cells can be reprogrammed to ectodermal induced neuronal (iN) cells by the overexpression of proneural transcription factor (TF) Ascl1 (Chanda et al., 2014; Vierbuchen et al., 2010). Co-expression of Brn2 and Mytl1 with Ascl1 (BAM), enhances reprogramming efficiency by suppressing a competing myogenic program (Chanda et al., 2014; Treutlein et al., 2016). During reprogramming, Ascl1 acts as a pioneer factor by binding to and opening closed chromatin regions, as well as modulating the binding of Brn2 (Wapinski et al., 2013), while Myt1l serves as multi-lineage repressor that maintains neuronal identity (Mall et al., 2017). The accumulation of abundant mCH is an epigenomic hallmark unique for mature neurons of mammalian brains, which could not be experimentally induced so far (Guo et al., 2014; Lister et al., 2013). mCH is recognized by MECP2 and serves critical gene regulatory functions in mature neurons (Chen et al., 2015; Gabel et al., 2015; Guo et al., 2014; Stroud et al., 2017). The reprogramming of fibroblasts to neurons using Ascl1 alone or in combination with Brn2 and Myt1l is a powerful way to interrogate the biochemical function of these TFs and given their prominent role during neural differentiation will also help dissecting their potential regulation of mC remodeling during neuron differentiation. Moreover, the role of DNA methylation in experimental induction of the neuronal lineage by TF reprogramming is not known yet and we currently lack a deep characterization of mC landscape of iN cells. Here we analyzed mC reconfiguration during iN cell reprogramming using whole-genome base-resolution methylome sequencing (MethylC-seq) and functionally perturbed methylation dynamics by genetic inhibition of Dnmt3a.

Base-resolution methylomes were generated for iN cells induced by either Ascl1 alone or all three TFs (BAM) in mouse embryonic fibroblasts (MEFs, Supplementary Table 1). We analyzed the methylomes of control MEFs, cells at initial (Ascl1 2d) and intermediate stages of reprogramming (Ascl1 5d and BAM 5d) and fully reprogrammed iN cells (Ascl1 22d and BAM 22d). 5d and 22d cells were isolated by fluorescence-activated cell sorting (FACS) using a neuronal TauEGFP reporter. We further generated methylomes for *in vitro* differentiation of neural progenitor cells (NPC, NPC 7d, NPC 14d and NPC 21d) under comparable culturing conditions. Remarkably, Ascl1 or BAM induced significant global accumulation of the neuronal-enriched non-conventional mCH methylation (Fig. 1a). mCH level reached an average of 0.28% in Ascl1 22d iN cells and a much greater level (0.57%) in BAM 22d iN cells, which is comparable to the frontal cortex tissue of 2 week old mice(Lister et al., 2013). In mature mouse cortical neurons (NeuN+), mCH is strongly enriched in the CAC context and mCAC accounted for approximately 34% of all mCH in mature neurons (Supplementary Fig. 1a). Similarly, we found neuronal reprogramming from fibroblasts was associated with a progressive establishment of mCAC predominance. In fully reprogrammed Ascl1 or BAM 22d iN cells, mCAC became the most abundant form (26% to 27%) of mCH. mCH is inversely correlated with gene expression in mature neurons(Guo et al., 2014; Lister et al., 2013). mCH was very weakly correlated to gene expression in MEFs or in intermediate reprogramming stages (BAM 5d) but became strongly inversely correlated with gene expression in BAM 22d cells suggesting a similar role of mCH in the brain and iN cells (Spearman r = −0.479, p-value < 2.2×10^−308^, Supplementary Fig. 1b). The accumulation of mCH in iN cells was correlated with a significant elevation of genome-wide mCG level from 72.7% in MEF to 79.5% in Ascl1 22d cells, and 82.5% in BAM 22d cells (Fig. 1a).

**Figure 1.**
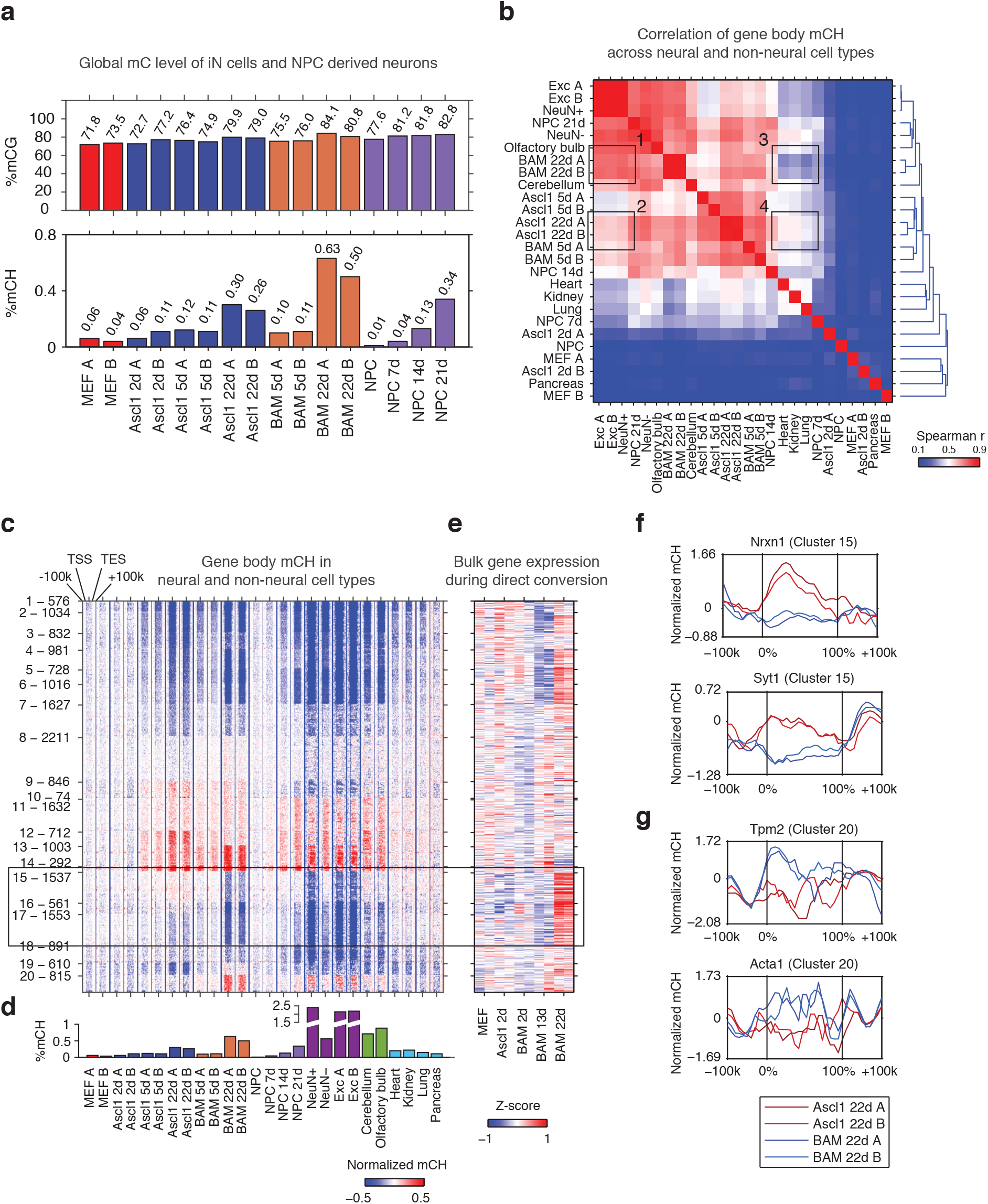
The Ascl1, Brn2 and Myt1l reprogramming factors induce an authentic, neuron-specific CH methylation pattern in fibroblasts. **a**, Global mCG (top panel) and mCH (bottom panel) levels of iN cells and NPC-derived cells. mCG and mCH accumulates in both maturing iN cells and differentiating NPCs **b**, Correlation of gene body mCH levels of iN cell reprogramming samples, NPC derived cells, neural and non-neural tissues. BAM 22d iN cells cluster most closely to mature cortical neurons. **c**, Left panel - K-means clustering of gene body mCH normalized to flanking regions (100 kb surrounding the gene). Reprogramming with BAM produces a global mCH profile more similar to cortical neurons compared to Ascl1 alone. NeuN+ and NeuN- indicate neuronal and glial nuclei separated using anti-NeuN antibody, respectively (Lister et al., 2013). Exc indicates purified nuclei from excitatory neuron expressing Camk2a+ (Mo et al., 2015). TSS - Transcription Start Site. TES - Transcription End Site. **d.** Global mCH level of samples compared in (c). **e.** Relative expression (Z-score) of bulk RNA-seq analysis of iN cell reprogramming. Gene expression is inversely correlated to gene body mCH. **f-g**, mCH pattern at genes with neuronal (**f**) and myogenic functions (**g**) in Ascl1 22d and BAM 22d iN cells. BAM 22d cells show greater depletion of mCH at synaptic genes and accumulation of mCH at myogenic genes, suggesting more efficient activation of neuronal genes and suppression of the alternate myogenic program compare to Ascl1 22d.

To evaluate whether iN cells recapitulate the mCH landscape of primary neurons in the brain, we compared iN cells to NPC-derived neurons of various times in culture, mature neurons freshly isolated from the brain, different brain regions as well as non-neural tissues with significant levels of mCH accumulation (global mCH/CH >= 0.1, Fig. 1b) (Hon et al., 2013; Shen et al., 2012). Using hierarchical clustering of gene body mCH levels, we found that gene body mCH in mature cortical neurons was more strongly correlated with BAM 22d cells (box 1) than with Ascl1 22d cells (box 2) reflecting the more mature state of BAM iN cells compared to Ascl1-iN cells(Treutlein et al., 2016). Accordingly, Ascl1 22d cells correlated more strongly to early reprogramming stages (Ascl1 5d and BAM 5d) and non-neural tissues (box 3,4 in Fig. 1b). Thus, as observed for gene expression patterns, BAM iN cells displayed a mCH landscape that is more similar to mature cortical neurons(Treutlein et al., 2016; Wapinski et al., 2013).

We identified genes showing similar or distinct mCH patterns between iN cells and mature neurons using K-means clustering (Fig. 1c,d and Supplementary Fig. 1c). We identified a large number (2,098, Cluster 15 and 16) of genes that were clearly hyper-methylated in MEFs, and immature iN cells but were strongly depleted of mCH in BAM 22d iN cells (box in Figure 1c). Notably, BAM 22d iN cells and matured cortical neurons showed similar hypo mCH for genes in Cluster 15 and 16. Through correlating with transcriptome analysis of iN cell reprogramming, we found that genes in Cluster 15 and 16 were actively expressed in successfully reprogrammed iN cells (Fig. 1e and Supplementary Fig. 1e)(Treutlein et al., 2016). Cluster 15 was enriched in gene functions such as synapse and metal ion binding (Supplementary Table 2). For example, *Nrxn1* and *Syt1* were depleted of mCH in BAM 22d cells but were enriched of mCH in immature neurons such as Ascl1 22d cells (Fig. 1f). We also found myocyte marker genes *Tpm2* and *Acta1* in Cluster 20, which shows greater level of mCH in BAM 22d iN than Ascl1 22d iN cells (Fig. 1g). This is consistent with our previous finding that Brn2 and Myt1l can suppress the cryptic myogenic program in iN cell reprogramming induced by Ascl1 (Treutlein et al., 2016). In summary, we found direct reprogramming using BAM factors produces a global mCH pattern more similar to cortical neurons, compared to using Ascl1 alone. mCH pattern in BAM iN cells is more permissive for the expression of neuronal and synaptic genes, and more repressive for the expression of the competing myogenic program.

To explore the role of mCH in regulating dynamic gene expression during reprogramming, we ranked genes by gene body mCH levels at an early stage of reprogramming (BAM 5d, Fig. 2a-c). Notably, genes with early mCH accumulation showed strongly reduced expression in the late stage of neuronal reprogramming (BAM 22d, Fig. 2b-c). We analyzed mCH accumulation at up- and down-regulated and static genes during reprogramming across a range of expression levels (Fig. 2d,e, Supplementary Fig. 2a, b). In all expression levels and reprogramming stages examined, down-regulated genes accumulated greater levels of mCH than genes with static or increased expression during reprogramming. Surprisingly, we found different patterns depending on the fold-change of gene expression: Mildly up-regulated genes accumulated intermediate mCH levels between down-regulated and static genes whereas strongly up-regulated genes and static genes were close to the MEF baseline of mCH levels (Fig. 2d,e). These results suggest a model that mCH is preferentially targeted to mildly dynamic genes and modulates their expression during reprogramming, for strongly differentially regulated genes, however, mCH is mostly accumulating at down-regulated genes.

**Figure 2.**
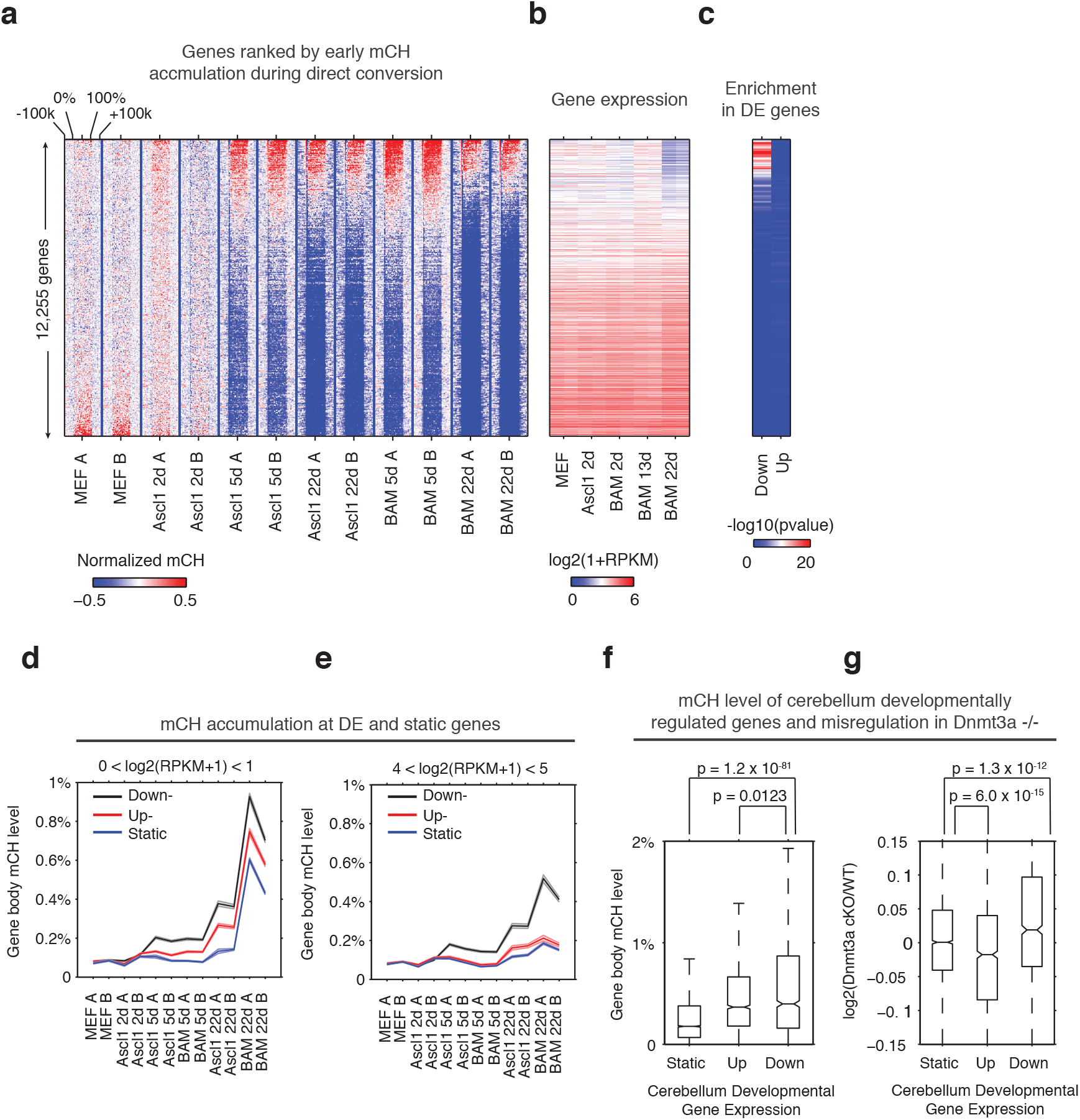
Early gene body mCH accumulation predicts later transcriptional down-regulation. **a-b**, Normalized gene body mCH (**a**) and transcript abundance (**b**) for genes ranked by early mCH accumulation at BAM 5d. Early mCH accumulation is strongly correlated to later gene repression. **c**, Significance (hypergeometric test) of the enrichment in down- and up-regulated genes. **d-e**, Gene body mCH dynamics of static, down- and up-regulated genes with different transcripts abundances - log2(RPKM+1) between 0 and 1 (**d**), between 4 and 5 (**e**) - during iN cell reprogramming. **f**, Gene body mCH level of cerebellum developmentally (P60 vs. P7) regulated genes that were actively expressed (RPKM>3). **g**, Alteration of cerebellum gene expression by Nes-Cre;Dnmt3a−/− for developmentally regulated genes.

To test whether mCH is associated with a similar effect on dynamic genes during mouse brain development, we analyzed mCH and gene expression in the cerebellum of mice with conditionally ablated *de novo* DNA methyltransferase Dnmt3a (Dnmt3a fl/fl;Nes-cre)(Frank et al., 2015; Gabel et al., 2015; Hon et al., 2013). Nes-cre;Dnmt3a fl/fl was shown to erase almost all mCH in mouse cerebellum(Gabel et al., 2015). Consistent with our observation in direct reprogramming, mCH level of genes down-regulated during cerebellum development (postnatal day 60 vs. day 7) was significantly greater than that of static (Wilcoxon rank sum test, p=1.2×10^−81^) or up-regulated genes (p=0.0123, Fig. 2f). Notably, the removal of mCH in Dnmt3a mutant brains did not affect the expression of developmentally static genes, but resulted in up-regulation of developmentally down-regulated genes (p = 1.3×10^−12^), and a down-regulation of developmentally up-regulated genes (p = 6×10^−15^, Fig. 2g). The result supported the role of *de novo* DNA methylation in modulating dynamically regulated genes during cell maturation.

Next, we wanted to determine the role of mCG dynamics in reprogramming. To examine the interaction between promoter (+/− 500bp from annotated transcription start sites (TSS)) mCG dynamics and gene expression, we ranked 7,427 differentially expressed (FDR<0.05, Fold change >= 2) genes by fold change between MEF and BAM 22d iN cells. We found a pronounced hyper CG methylation signature at the promoter of genes that become repressed during reprogramming (Fig. 3b,c), such as the *Col1a2* locus (Fig. 3a). Moreover, *Col1a2* TSS is heavily methylated in NPC and NPC derived neurons, but not in *in vivo* grown cells such as fetal cortex or mature cortical neurons (NeuN+) or glia (NeuN-), suggesting direct reprogramming remodeled the methylation state of *Col1a2* promoter to that reminiscent *in vitro* grown neurons (Fig. 3a). To compare promoter mCG between direct reprogrammed iN cells, *in vitro* and *in vivo* grown cells, we identified 4,496 promoters that became significantly hyper CG methylated in BAM 22d iN cells compared to MEFs (Supplementary Fig. 3a and Table 3). Promoters that became hyper CG methylated during direct reprogramming showed significantly greater mCG in BAM 22d iN cells than in *in vivo* grown cells (p<2.9×10^−7^, rank sum test, Supplementary Fig. 3b), suggesting promoter hyper CG methylation is an epigenomic signature of direct reprogramming. During reprogramming, mCG at promoters showed a strong inverse correlation with gene expression (Fig. 3d). This negative correlation between promoter mCG and gene expression in iN cells attenuates rapidly within 2kb of the TSS, which is strikingly different from primary human or mouse tissues (Fig. 3e and Supplementary Fig. 3c). Pairwise comparisons of brain cortex to kidney, cerebellum, heart and lung supported the previous analysis of human tissues that the peak negative correlation between mCG and gene expression was found at 2-3 kb downstream of TSS (Fig. 3e and Supplementary 3c) (Schultz et al., 2015). Across a broad range gene expression levels, we could identify a subset of genes clearly showing down-regulation and promoter hyper CG methylation in BAM 22d iN cells compared to MEF (Supplementary Fig. 3d). This subset of genes were also more actively expressed in MEF than in adult mouse cortex (Fig. 3f,g). Therefore the downregulation of genes with hyper CG methylated promoter during direct programming is consistent with their repressed expression in *in vivo* grown neurons. Consistent with their active expressions in MEF, genes with promoter hyper CG methylation induced during reprogramming were strongly enriched in fibroblast specific gene ontology terms such as extracellular matrix or collagen (Fig. 3h).

**Figure 3.**
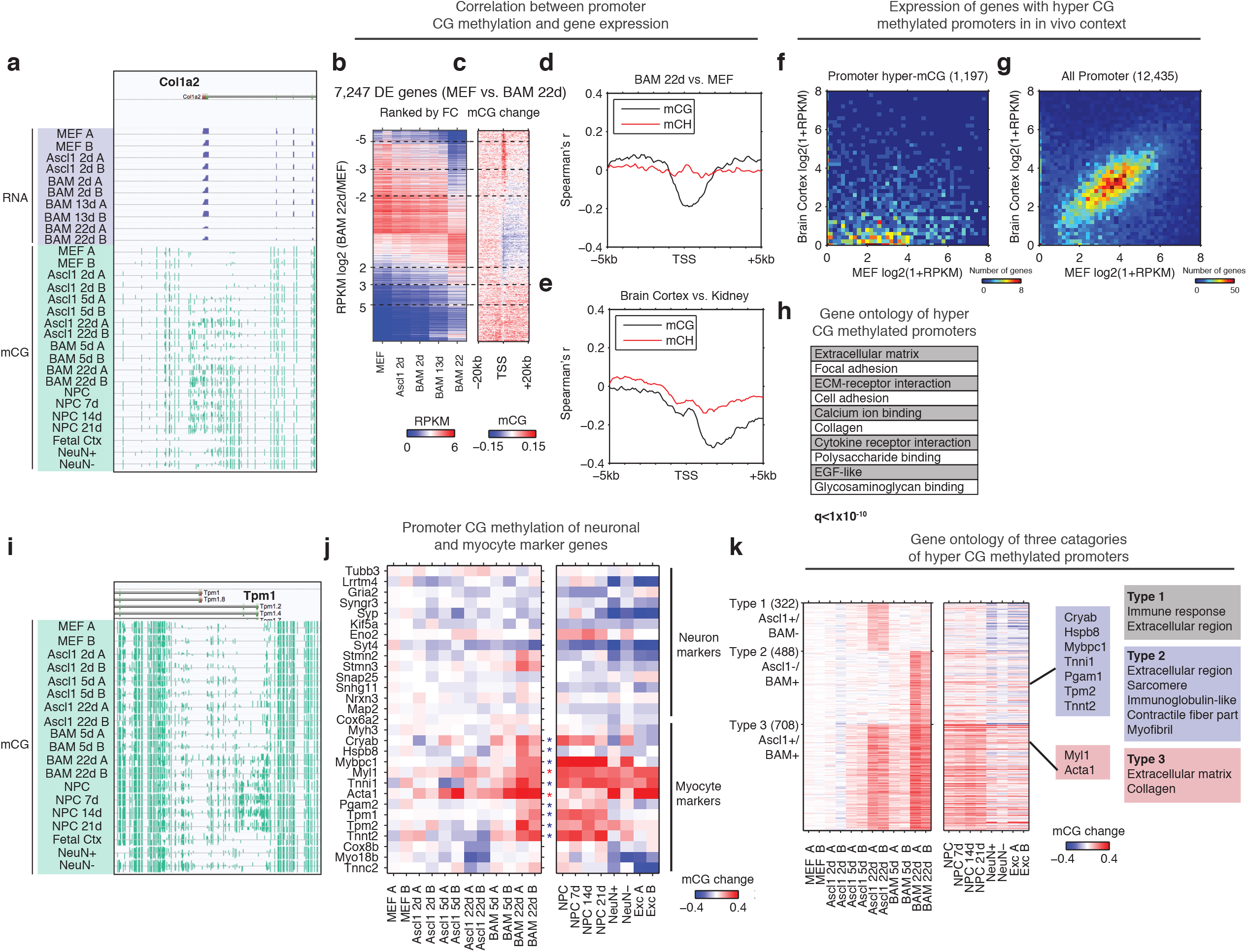
Silencing of key non-neuronal programs is associated with promoter hyper CG methylation. **a**, Browser view of fibroblast marker gene *Col1a2* promoter methylation dynamics and corresponding gene expression during reprogramming. The promoter is strongly methylated throughout NPC differentiation, but is unmethylated in fetal cortex, cortical neurons and glia. However, it gradually accumulates mCG through the course of reprogramming for both Ascl1 alone and BAM factors. **b**, Differentially expressed genes between BAM 22d iN and MEF were ranked by their relative fold-changes. **c**, mCG change between MEF and BAM 22d in +/− 20kb regions surrounding TSS of differentially expressed genes. Hyper CG methylation was found at TSS for down-regulated genes. **d**, Correlation between mCG change and differential gene expression between BAM 22d iN and MEF in +/− 5kb region surrounding TSS. **e**, Correlation between mCG change and differential gene expression between brain cortex and kidney in +/− 5kb regions surrounding TSS. **f-g**, Scatter plots comparing the expression of genes with hyper CG methylation promoter in BAM 22d iN cells (**f**) or all genes (**g**) between MEF and brain cortex. Hyper CG methylation genes in BAM 22d cells are strongly repressed compared to all genes in the *in vivo* context. **h**, Gene ontology term enrichment of genes with hyper CG methylation promoters in BAM 22d iN cells. **i**, Browser view of promoter mCG dynamics of myocyte marker gene *Tpm1*. The promoter is already methylated through NPC differentiation, but is unmethylated in fetal cortex and cortical neurons and glia. However, it remains unmethylated when reprogramming with Ascl1 alone, and only accumulates mCG in the presence of Brn2 and Myt1l in BAM. **j**, Promoter mCG dynamics of neuron and myocyte marker genes during reprogramming. Blue asterisks indicate promoters hyper-methylated in BAM 22d iN, but not in Ascl1 22d iN. Red asterisks indicate promoters hyper-methylated in both BAM 22d iN and Ascl1 22d iN. A majority of myocyte genes are only hyper methylated by BAM and not Ascl1 alone. **k**, Left: Heatmaps showing promoter mCG during neuronal reprogramming, NPC differentiation and in primary neurons. Promoters were categorized into three groups showing hyper-mCG (mCG increase > 0.1) in Ascl1 22d only, BAM 22d only or both type of iN cells. Right: Gene ontology enrichment of promoters for the 3 groups. Promoters of fibroblast genes were hyper CG methylated in both Ascl1 alone and BAM iN cells, but only BAM 22d cells exhibit strong hyper CG methylation at myogenic genes.

We further asked whether the suppression of myogenic program by BAM factors involves the repression of key gene expression by DNA methylation. Indeed, promoters of a significant fraction (8/15) of analyzed myocytic marker genes (*Cryab*, *Hspb8*, *Mybpc1*, *Tnni1*, *Pgam2*, *Tpm1*, *Tpm2* and *Tnnt2*) were hyper-methylated in BAM 22d and NPC derived neurons but not in Ascl1 22d iN (Fig. 3i,j) (Treutlein et al., 2016). This result correlates with previous data showing repressed myocyte-specific genes in BAM iN cells (Treutlein et al., 2016). Only a subset of myocytic marker genes (*Myl1*, *Tnni1* and *Acta1*) were also hyper CG methylated in cortical neurons, suggesting the promoter hyper-methylation at myocytic genes is in part an epigenetic program specific for *in vitro* grown neurons. We further grouped promoters into three types based on their methylation states in Ascl1 22d or BAM 22d iN cells (Fig. 3k). Type 2 promoters that were specifically hyper CG methylated in BAM 22d cells but not Ascl1 22d, and were enriched in skeletal muscle related function such as sarcomere, contractile fiber part and myofibril (Fig. 3k). Remarkably, promoters showing hyper CG methylation during reprogramming were marked with elevated H3K27me3 levels in cortical neurons (p<1.2×10^−50^, rank sum test, Supplementary Fig. 3e), suggesting these loci are repressed with polycomb repressive complexes (PRC) in *in vivo* grown neurons. Promoter hyper CG methylation of fibroblast and myocyte specific genes may reflect an alternative repressive mechanism (DNA methylation vs. PRC) being utilized during direct reprogramming or cell culture. Taken together, the suppression of fibroblast and myogenic programs during reprogramming was associated with promoter hyper CG methylation, which indicates the potential role of DNA methylation in mediating the suppression of historical and competing programs during reprogramming.

Discrete regions with low levels of DNA methylation have previously been found to indicate TF binding (Mo et al., 2015; Schultz et al., 2015; Stadler et al., 2011). To explore the interaction between CG methylation and TF binding, in particular the binding of ASCL1, BRN2 and MYT1L, we identified 10,075 and 15,093 differentially methylated sites (DMSs) showing reduced mCG during Ascl1-only and BAM induced reprogramming, respectively (Fig. 4a,c and Supplementary Table 3). 96% of Ascl1-only DMSs and 95.7% of BAM DMSs were located more than 2.5kb away from the closest TSS. Thus the vastly majority of DMSs are likely to be associated with distal gene regulatory activities. DMSs were grouped based upon their demethylation kinetics (Fig. 4a,c). Among the sites that become demethylated in Ascl1-only reprogramming we found a striking co-enrichment of ASCL1 binding, in particular at early demethylating sites (Fig. 4b and Supplementary Fig. 4c), suggesting that ASCL1 can recruit a DNA demethylation machinery to its target sites. This fits well with Ascl1’s known function as a strong transcriptional activator (Castro et al., 2011; Raposo et al., 2015; Wapinski et al., 2013). Surprisingly, however, only 0.6% (99/15,093) BAM DMSs were overlapped with ASCL1 peaks (Fig. 4d and Supplementary Fig. 4c). DMSs demethylated in early stage of BAM induced reprogramming (BAM DMS group 3) are enriched in BRN2 binding sites in MEF (Fig. 4d). BAM DMSs demethylated in fully reprogrammed BAM 22d iN cells showed a moderate overlap with BRN2 binding sites in neural stem cells (NSC, Fig. 4d). BRN2 binding is known to be dependent on existing open chromatin and pioneer factors such as ASCL1 (Wapinski et al., 2013). This result suggests BRN2 is able to at least partially access its physiological binding sites in fully reprogrammed BAM 22d iN cells. Although Myt1l serves as multi-lineage repressor during reprogramming, we found only 0.3% (38/15,093) BAM DMSs overlapped with MYT1L binding sites (Supplementary Fig. 4a-c). Therefore MYT1L binding does not majorly contribute to mCG remodeling during the direct reprogramming.

**Figure 4.**
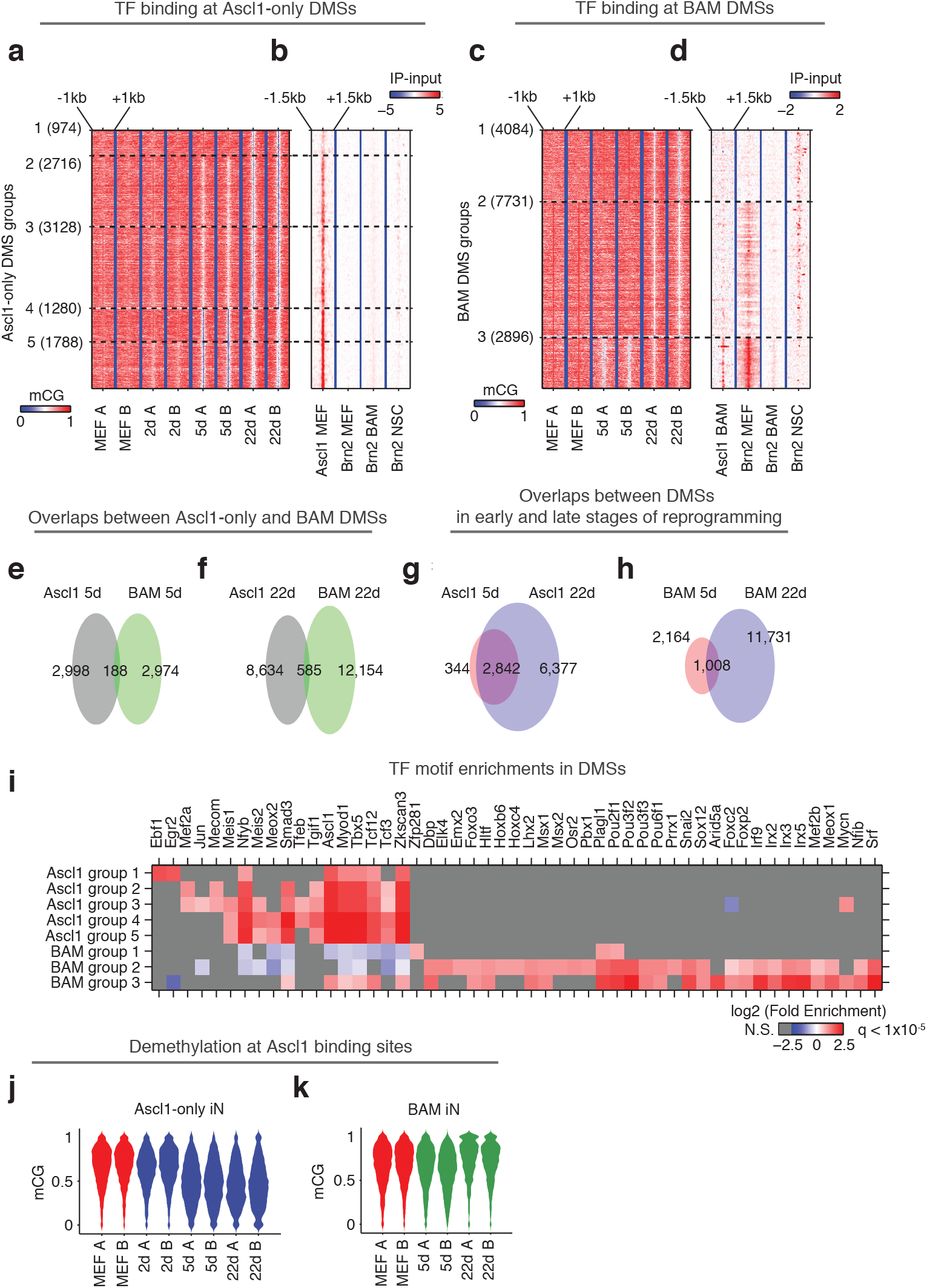
Ascl1 and BAM factors induce distinct CG methylation reconfigurations. **a,c**, DMSs identified during reprogramming driven by Ascl1 (**a**) or BAM (**c**) expression was ordered by the kinetics of mCG remodeling. **b,d**, Signal intensity of Ascl1 and Brn2 ChIP-seq were plotted for 3 kb regions surrounding the DMSs. **e-h**, The overlaps of DMSs identified at different stages of reprogramming driven by Ascl1 or BAM expression. **i**, TF binding motif enrichment in DMS groups shown in (**a**) and (**c**). Insignificant enrichments (q > 1×10^−5^) were shown as gray. j, The distribution of mCG level at Ascl1 peaks during Ascl1-only induced cells during reprogramming. **k**, The distribution of mCG level at Ascl1 peaks in BAM induced cells during reprogramming.

Consistent with the drastically different representations of ASCL1 binding sites in Ascl1-only DMSs and BAM DMSs, less than 10% DMSs were shared between Ascl1-only and BAM induced reprogramming (Fig. 4e-f). In contrast, most DMSs found in Ascl1 5d (89.2%) were also found in Ascl1 22d (Fig. 4g), suggesting the little overlap between DMSs demethylated during Ascl1-only and BAM induced reprogramming reflects true biological differences. To further illuminate the nature of the demethylating regions we performed motif enrichment analysis. As expected, the Ascl1 binding motif was strongly enriched in Ascl1-only DMSs (Fig. 4i). The DMSs found in the two types of iN cell reprogramming were enriched in completely different TF binding motifs (Fig. 4i). Accordingly, the Ascl1 motif was only partly enriched in BAM DMSs demethylated in early reprogramming (BAM DMS group 3), but was depleted in BAM DMSs emerged in later stages of reprogramming (BAM DMS group 1). As expected, the Brn2 (Pou3f2) motif along with several other Pou-Homeodomain motifs were enriched in BAM DMSs but not in any Ascl1-only DMSs. Moreover, we found enrichment of additional motifs of TFs many of which with prominent neuronal function such as Lhx2, Emx2, Nfib and Mef2 in BAM DMSs. These findings indicate that the dynamic DMSs during iN cell reprogramming are enriched in DNA regulatory elements and their proper activation involves DNA demethylation as part of the chromatin remodeling of a fibroblast to a neuronal configuration.

Given the striking enrichment of ASCL1 binding at Ascl1-only DMSs, we next analyzed the mCG dynamics of just the Ascl1 binding sites during reprogramming. Although only a moderate 28.4% (935/3296) of Ascl1 binding sites (not already showing lowly methylation in MEF) were overlapped with Ascl1-only DMSs, our quantitative analysis found that Ascl1 expression in fibroblasts induced a substantial DNA demethylation at the vast majority of Ascl1 binding sites in early reprogramming Ascl1 5d with persistence of reduced mCG levels to 22 days (Fig. 4j). During BAM-induced reprogramming, the ASCL1-bound sites behaved completely differently. Early in the reprogramming process Ascl1 did induce demethylation at many of its target sites, but to a much more moderate degree (BAM 5d, Fig. 4k). Surprisingly, these changes were only transient, as the demethylation of Ascl1 binding sites was reversed in the mature stage of BAM iN reprogramming (BAM 22d). The results more directly suggest that Brn2 and Myt1l functionally interfere with Ascl1-mediated local demethylation. Since ASCL1 chromatin binding is unaffected by other transcription factors, this observation indicates that addition of Brn2 and Myt1l modifies ASCL1’s ability to induce DNA demethylation at its chromatin binding sites and that the two reprogramming strategies are associated with drastically different DNA methylation remodeling (Fig. 4j,k). Such modulation of ASCL1 by BRN2 is supported by the co-binding of the two TFs at about about a quarter of ASCL1 sites in MEF (Wapinski et al., 2013).

Next, we assessed the expression of known DNA methylation regulators throughout the entire reprogramming process. We found a strong up-regulation of the *de novo* DNA methyltransferase Dnmt3a, the tet methylcytosine dioxygenase 3 (Tet3) involved in DNA demethylation and the mCH reader Mecp2 expressions during reprogramming (Fig. 5a)(Chen et al., 2015; Gabel et al., 2015; Guo et al., 2014). The expression of Uhrf1 that encodes a SRA domain protein required for DNMT1-mediated mCG was drastically reduced in BAM 13d and 22d (Fig. 5a)(Bostick et al., 2007), suggesting that the maintenance DNA methylation activity became repressed in post-mitotic iN cells. The other DNA methylation regulators were not dynamically regulating during the reprogramming process. Of the regulated genes, we found that ASCL1 directly binds to Dnmt3a promoter. ASCL1 occupies a binding site 1,957 bp downstream of Dnmt3a transcription start site (TSS) in MEF and initial reprogrammed cells suggesting a direct contribution of Ascl1 to the upregulation of Dnmt3 during iN cell reprogramming (Fig. 5b).

**Figure 5.**
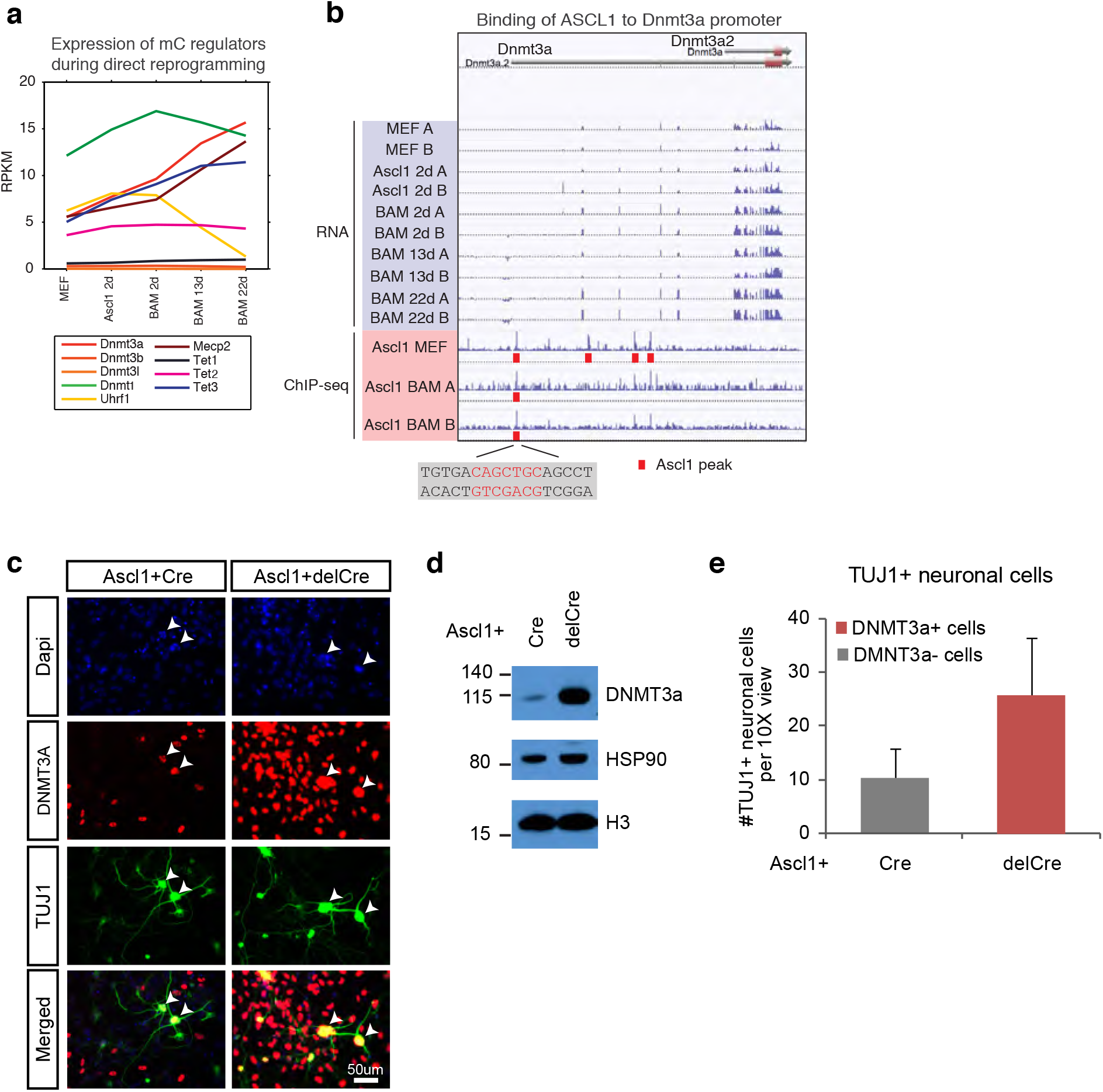
*de novo* DNA methylation is required for efficient reprogramming from fibroblasts to neurons. **a**, Expression of DNA methylation regulators and readers during neuronal reprogramming. **b**, Ascl1 ChIP-seq peak is located proximal to Dnmt3a promoter. **c**, Immunostaining of Dnmt3a/3b^fl/fl^ cells 14d post-Ascl1 induction with dox. There appears to be a similar reprogramming efficiency when comparing the CreGfp (left) and delCreGfp (right) conditions. However, most TUJ1+ iN cells in the CreGfp condition were still expressing low levels of DNMT3A (white arrows) despite co-infection with CreGfp. **d**, Western blot showing efficient knock-out of DNMT3A in Ascl1+Cre expressing cells compared to Ascl1+delCre control 13 days post induction of Ascl1. **e**, Average counts of TUJ1+ iN cells per 10X field of view 14d post-Ascl1 induction that are co-infected with 1) CreGfp and Dnmt3a- (gray) or 2) delCreGfp and Dnmt3a+ (red) (error bars are stderr, n=3).

To functionally query the role of *de novo* DNA methylation during reprogramming, we then acutely ablated the two main *de novo* DNA methylases by co-infecting fibroblasts isolated from Dnmt3a/3b^fl/fl^ mice with lentivirus expressing intact Cre recombinase (Cre) or a non-functional truncated form of Cre recombinase (delCre). To initiate reprogramming we co-infected the cells with a doxycycline (dox)-inducible TetO-Ascl1 lentivirus. Dox was added 2-4 days post-infection to allow for degradation of Dnmt3a mRNA and protein before inducing Ascl1 expression for direct conversion of MEFs to neurons. Dnmt3b is not expressed through the course of reprogramming (Fig. 5a). Cells were fixed and immunostained with the neuronal marker TUJ1 and DNMT3A (Fig. 5c, Supplementary Fig. 5a) 14 days after Ascl1 induction. Despite efficient Dnmt3a depletion in the total fibroblast population (Fig. 5d, Supplementary Fig. 5c) we observed a large fraction of Dnmt3a-positive iN cells in the Cre-infected group, presumably due to incomplete degradation of Dnmt3a or escape from Cre recombination, that reprogram into neurons (Fig. 5c, Supplementary Fig. 5d-f). We therefore considered only the Dnmt3a-negative iN cells in the Cre-treated group. As shown in Fig. 5f, the reprogramming efficiency of Dnmt3a-negative cells (gray) was reduced compared to control-infected, Dnmt3-positive cells (red), indicating that de novo DNA methylation plays a role in efficient iN cell generation from fibroblasts.

Our study found that fully reprogrammed iN cells accumulate abundant mCH, which is an epigenomic signature of post-mitotic neurons in the mammalian brain (Guo et al., 2014; Lister et al., 2013). To our knowledge, high level (e.g. mCH/CH > 0.1%) has not been previously reported in any *in vitro* neuronal differentiation or direct reprogramming models. Robust mCH accumulation suggests that iN cells recapitulates a major epigenomic signature that starts to emerge during the second postnatal week of mouse brain development and supports the notion that fully reprogrammed iN cells resemble differentiated neurons (Lister et al., 2013; Vierbuchen et al., 2010). The observation that BAM induced iN cells further acquired a global mCH landscape reminiscent to mature neurons is consistent with previous findings that BRN2 and MYTL1 can enhance neuronal maturation during reprogramming (Chanda et al., 2014). The reprogramming of fibroblasts to neurons and induction of mCH by overexpressing merely 1 or 3 TFs provides a tractable system to study the dynamic regulation of genome-wide DNA methylation. For example, mCH accumulation during reprogramming is associated with the upregulation of Dnmt3a and may be regulated by the direct binding of ASCL1 to Dnmt3a promoter. In addition, inducible expression of Ascl1 allowed us to conclude that ASCL1 induces local demethylation at most of its binding sites.

mCH is inversely correlated with gene expression in the brain and has been shown to mediate gene repression through MECP2 (Chen et al., 2015; Gabel et al., 2015; Lister et al., 2013; Stroud et al., 2017). Although the timing of mCH accumulation parallels neuron maturation both *in vivo* during *in vitro* reprogramming, little is known whether mCH plays any role in regulating brain development. Our integrative analysis of transcriptomic and DNA methylome datasets for neuronal reprogramming and published work on cerebellar development shows that mCH is enriched in genes that are downregulated during development (Frank et al., 2015), suggesting that mCH plays a role in gene repression during brain development. Reanalysis of the Dnmt3a conditional knock-out transcriptome data supported our model - developmental downregulated genes become activated in the absence of mCH (Gabel et al., 2015). In both mouse and human, mCH accumulation start around birth and plateaus at early adolescence. Our results collectively suggest that mCH facilitates neuronal maturation during development through gene expression.

Direct conversion from fibroblasts to neurons is associated with distinct promoter hypermethylation, which is unique to iN cells and NPC differentiated cells and is not observed during *in vivo* neuron differentiation. We thus found iN cells exhibit both consistent (e.g. mCH) and different (e.g. promoter methylation) epigenomic features compared to mature cortical neurons. Our study suggests that *de novo* DNA methylation contributes to cellular reprogramming by suppressing the donor fibroblast and the competing myogenic programs. The promoters targeted by hyper methylation is an independent set from ones bound by multi-lineage suppressor Mylt1l. In summary, our genomic analysis suggests both types of DNA methylation - mCH and mCG serve repressive functions during cellular reprogramming and primarily target gene bodies and promoters, respectively. This hypothesis is supported by our functional experiment showing that fibroblast cells with ablated Dnmt3a locus have reduced reprogramming potential, which supports hypothesis that *de novo* DNA methylation is necessary for efficient reprogramming.

## Acknowledgement

The work is supported by the Howard Hughes Medical Institute (J.R.E), NIH P50-HG007735 (H.Y.C), CIRM RB5-07466 (H.Y.C., M.W.) and NIH R01 DK092883 (M.A.G).

## Methods

### Cell derivation

Homozygous TauEGFP knock-in mice were bred with C57BL/6 mice (Jackson Labs) to generate heterozygous embryos(Tucker et al., 2001). The derivation of fibroblast cultures (TauEGF MEFs) was performed by isolating only the limbs from E13.5 embryos, which were then chopped into small pieces, trypsinized for 15min at 37°C and plated in MEF media, containing 10% cosmic calf serum (Hyclone), 0.008% Beta mercaptoethanol (Sigma), MEM non-essential amino acids, Sodium Pyruvate, and Penicillin/Streptomycin (Pen/Strep) (all from Invitrogen). Neural progenitor cells (TauEGFP NPCs) were derived by harvesting the two cortical lobes from the brain of E12.5 embryos, which are then incubated in N3 media containing DMEM/F12, N2 supplement, Pen/Strep (all from Invitrogen) and 20ug/ml Insulin (Sigma) at 37°C for 10min. The cortical tissue was then gently dissociated with a 1mL micro-pipette, passed through a 0.7um filter before being plated in N3 media with EGF (20ng/mL) and FGF (10ng/mL) onto a cell culture dish that was previously coated with polyornithine (PO) (Sigma P3655) for at least 4hrs followed by laminin (LAM) (Sigma L2020) overnight.

For homozygous Dnmt3a/3b^fl/fl^ MEFs, Dnmt3a^fl/fl^ and Dnmt3b^fl/fl^ mice on a C57BL/6 background were crossed to generate Dnmt3a/3b^fl/fl^ mice(Challen et al., 2014; Dodge et al., 2005; Kaneda et al., 2004). Whole body MEFs were harvested from E13.5 embryos by removing the head, spinal cord and red organs, minced into small pieces, trypsinized and plated in MEF media on gelatinized flasks to derive fibroblast cultures(Jozefczuk et al., 2012). Both MEFs were expanded for 3 passages, while NPCs were expanded 3 to 4 passages prior to experiments.

### Direct conversion of fibroblast to neuron

Lentivirus was produced, and TauEGFP MEFs were co-infected with reverse tetracyclin transactivator (rtTA) and doxycycline (dox) inducible TetO-Ascl1 alone or with TetO-Brn2 and TetO-Mytl1 as previously described(Marro and Yang, 2014). Dox was added with fresh MEF media 16-20hrs post lentiviral infection, and cultures were switched to N3+B27 media containing DMEM/F12, N2 and B27 supplements, Pen/Strep (all from Invitrogen) and 20ug/ml Insulin with dox after 2 days. Cells were harvested at 48 hours, 5d and 22d post-dox induction for MethylC-seq. Control MEFs were not infected and harvested 48hr after addition of dox. For 5d and 22d, cells were FAC (Fluorescence-activated cell)-sorted for TauEGFP+ cells to select for cells that were reprogramming. To ensure that reprogramming efficiencies are comparable, immunofluorescence staining for Tuj1 was performed for each batch of cells at day 14, and only samples that average at least 20 Tuj1+ neurons per 10X field of view were used. In addition for FAC-sorted samples, only samples containing >5% TauEGFP+ cells was used.

Dnmt3a/3b^fl/fl^ MEFs were co-infected with rtTA, TetO-Ascl1 and either CreGFP or delCreGFP (both with constitutive FUW promoters). DelCreGFP contains a truncated and non-functional version of Cre and was used as a control. Fresh MEF media was added 16-20hr after infection. Dox was then added between 2-4 days later to allow knockout of the Dnmt3a/b locus by Cre and degradation of the remaining protein before reprogramming with Ascl1.

### *In-vitro* differentiation of neural progenitor cells into neurons

P3-4 NPCs were seeded into cell culture plates that were previously coated with PO+LAM in N3+EGF+FGF media. After one day, the media was replaced with fresh differentiation media containing N2+B27 media, without EGF and FGF. Half media replacement with fresh N2+B27 media was performed every alternate day. Control NPCs were harvested for MethylC-seq 2 days after seeding (without addition of differentiation media). Differentiating cells were harvested for MethylC-seq at 7d, 14d and 21d after addition of differentiation media and were FAC-sorted for TauEGFP+ cells to select for neuronal cells.

### DNA extraction for MethylC-seq

DNA extraction was performed using the Qiagen DNeasy Blood and Tissue kit (#69506) with some modifications to the protocol. Fresh cells were washed once with PBS, then re-suspended in 900uL ATL, 100uL Proteinase K, 20uL of RNAse A (Invitrogen 12091021) and 200uL AL. Cell suspension is then incubated at 56°C for 10min. Then 1mL of 100% Ethanol is added and mixture was briefly vortexed before loading onto a spin-column. DNA is then washed on the column following Qiagen’s protocol and finally eluted into 50-100uL AE.

### Immunofluorescence and cell counting

Cells were fixed and counted as previously described(Wapinski et al., 2017). Briefly, cells were fixed with 4% paraformaldehyde at 14d post-dox, blocked with 5% cosmic calf serum (CCS), incubated with primary antibodies for at least an hour, followed by secondary antibodies for at least half an hour. Ten 10X images were then taken per biological replicate (MEFs derived from different embryos), and the relevant cells were counted. Tuj1+ neuronal cells were counted manually, and only cells with neurite extensions that are at least three times the length of the cell body diameter are included in the counts. ImageJ was used for counting DAPI, Dnmt3a+ and Cre/delCreGfp+ cells (Schneider et al., 2012). A threshold was first set to eliminate background noise and kept consistent for all images within a single biological replicate. Then “Watershed” was ran to distinguish distinct nuclei, and “Analyze particles” was used (size=0.01-Infinity) to count the number of cells. For each replicate, the number of cells per 10X view was taken as an average of all the images taken.

### Antibodies

Rabbit anti-Tubb3 (Tuj1, Covance MRB-435P), rabbit anti-Dnmt3a (H-295, Santa Cruz Biotech sc-20703), chicken anti-GFP (Abcam ab13970). Secondary Alexa-conjugated antibodies were used at 1:1000 (all from Invitrogen).

### MethylC-seq

MethylC-seq libraries were prepared as previously described(Urich et al., 2015), except regular lambda DNA (Promega cat. # D1501) isolated from *dcm*+ E.Coli was spiked into samples as the control for non-conversion rate. MethylC-seq libraries were sequenced on Illumina HiSeq 2500. Non-conversion rate was computed from each sample after excluding CAG and CTG trinucleotides from lambda DNA sequence. MethylC-seq reads were mapped to mm9 reference genome using MethylPy (https://bitbucket.org/schultzmattd/methylpy/wiki/Home)(Lister et al., 2013; Ma et al., 2014). To ensure the correct calling of mCG and mCH, we removed cytosine positions with potential single nucleotide variants located at cytosines or immediately downstream of cytosines as previously described(Luo et al., 2016).

To compare the methylome of iN cells to primary neurons and other mouse tissues, methylome reads of NeuN+ neurons (GSM1173786), NeuN-glia (GSM1173787), Camk2a+ excitatory neurons (GSM1541958 and GSM1541959), cortex (GSM1051153), cerebellum (GSM1051151), olfactory bulb (GSM1051159), heart (GSM1051154), kidney (GSM1051156), lung (GSM1051158) and pancreas (GSM1051160) were downloaded from NCBI SRA and processed identically as iN cell methylome data.

Differentially methylated sites were identified between pairs of iN cell samples using DSS with FDR < 0.1(Feng et al., 2014).

### ChIP-seq data

Ascl1 and Brn2 ChIP-seq dataset for iN cell samples were previously published in GSE43916 (Wapinski et al., 2013) - Ascl1 MEF (SRR935631, SRR935632), Ascl1 BAM (SRR935633, SRR935635, SRR935634), Brn2 MEF (SRR935643, SRR935644), Brn2 BAM (SRR935645, SRR935646), Brn2 NSC (SRR935647, SRR935648). Sequencing reads were mapped to mm9 reference genome using bowtie2 2.1.0(Langmead and Salzberg, 2012). ChIP-seq peaks were identified using MACS2 2.0.10 with q value < 0.01. Peaks called from the two replicates of Brn2 BAM ChIP-seq were combined. To analyze DNA demethylation induced by TF bindings, Ascl1 and Brn2 peaks that overlapped with lowly methylate regions (UMRs or LMRs) identified from Ascl1 2d methylome were removed. UMRs and LMRs were identified using MethylSeekR with m=0.5 and 5% FDR(Burger et al., 2013).

### RNA-seq data

For all reprocessing of published RNA-seq data, RNA-seq reads were downloaded from NCBI SRA and mapped to mm9 Refseq gene annotation using STAR 2.4.0(Dobin et al., 2013). Mapped RNA-seq reads were counted and summarized to the gene level with HTSeq 0.6.1 followed by normalization and computation of RPKM using edgeR 3.8.6 (Anders et al., 2015; Robinson et al., 2010). Differentially expressed genes were identified using edgeR 3.8.6 with FDR < 0.05 and Fold Change >= 2(Robinson et al., 2010).

RNA-seq dataset of iN cell reprogramming was downloaded from GSE43916(Wapinski et al., 2013). Single cell gene expression dataset of iN cell reprogramming was previously published(Treutlein et al., 2016). Mouse tissue RNA-seq dataset was downloaded from NCBI GEO accessions cerebellum (GSM723768), cortex (GSM723769), heart (GSM723770), kidney (GSM723771), lung (GSM723773) and olfactory bulb (GSM850911)(Shen et al., 2012). Cerebellum developmental RNA-seq dataset was downloaded from NCBI SRA accessions P7 (SRR1557065, SRR1557066, SRR1557067), P14 (SRR1557068, SRR1557069, SRR1557070) and P60 (SRR1557071, SRR1557072, SRR1557073). Dnmt3a −/−;Nes-cre gene expression dataset was downloaded from NCBI GEO accessions WT (GSM1464557, GSM1464558, GSM1464559) and Dnmt3a KO (GSM1464560, GSM1464561, GSM1464562).

### Accession numbers

Methylome profiles are visualized using an AnnoJ browser at http://neomorph.salk.edu/iN_transdifferentiation.php.

## Supplementary Figure Legends

**Supplementary Figure 1.**
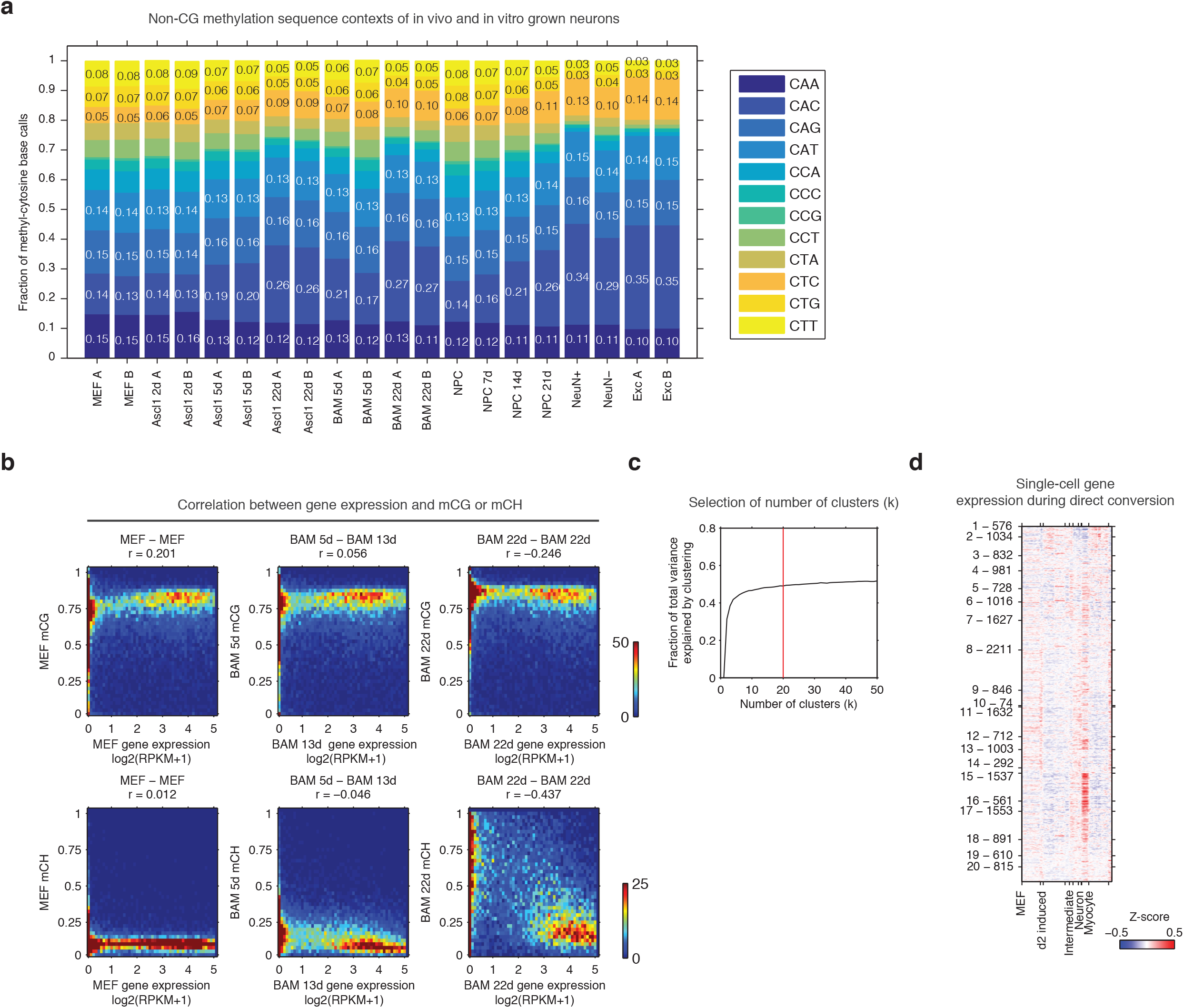
**a**, The frequency of tri-nucleotide contexts in mCH base calls for iN cells and cortical neuron samples. **b**, Correlations between iN cell gene body mCG or mCH levels and gene expressions. **c**, The fraction of total variance explained plotted as a function of the number of clusters for the kmeans clustering shown in Fig. 1c. **d**, Single cell gene expressions plotted for gene clusters shown in Fig. 1c.

**Supplementary Figure 2.**
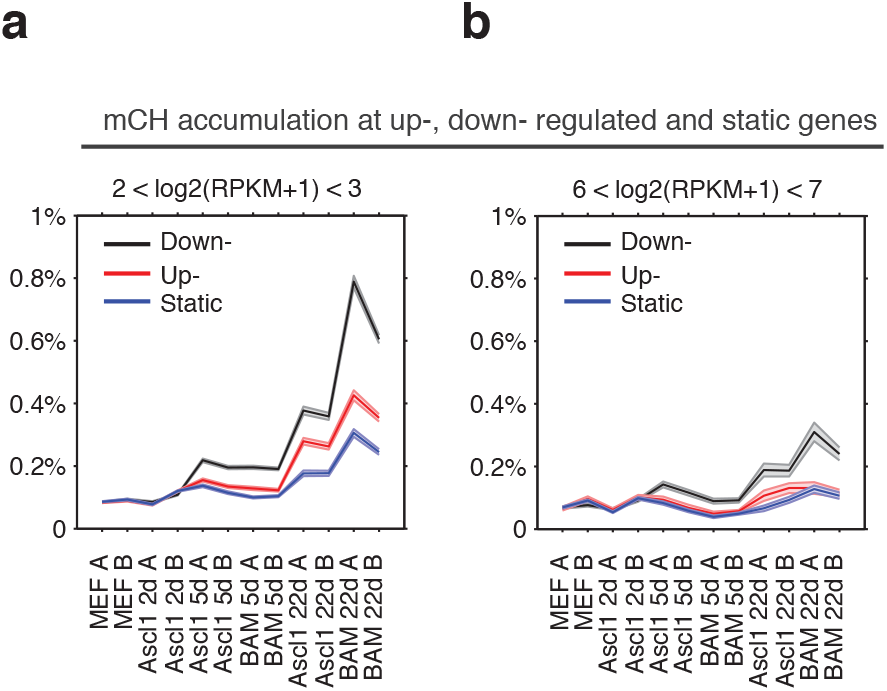
**a-b,** Gene body mCH dynamics of developmentally static, down- and up-regulated genes with different transcripts abundances - log2(RPKM+1) between 2 and 3 (**a**), between 6 and 7 (**b**).

**Supplementary Figure 3.**
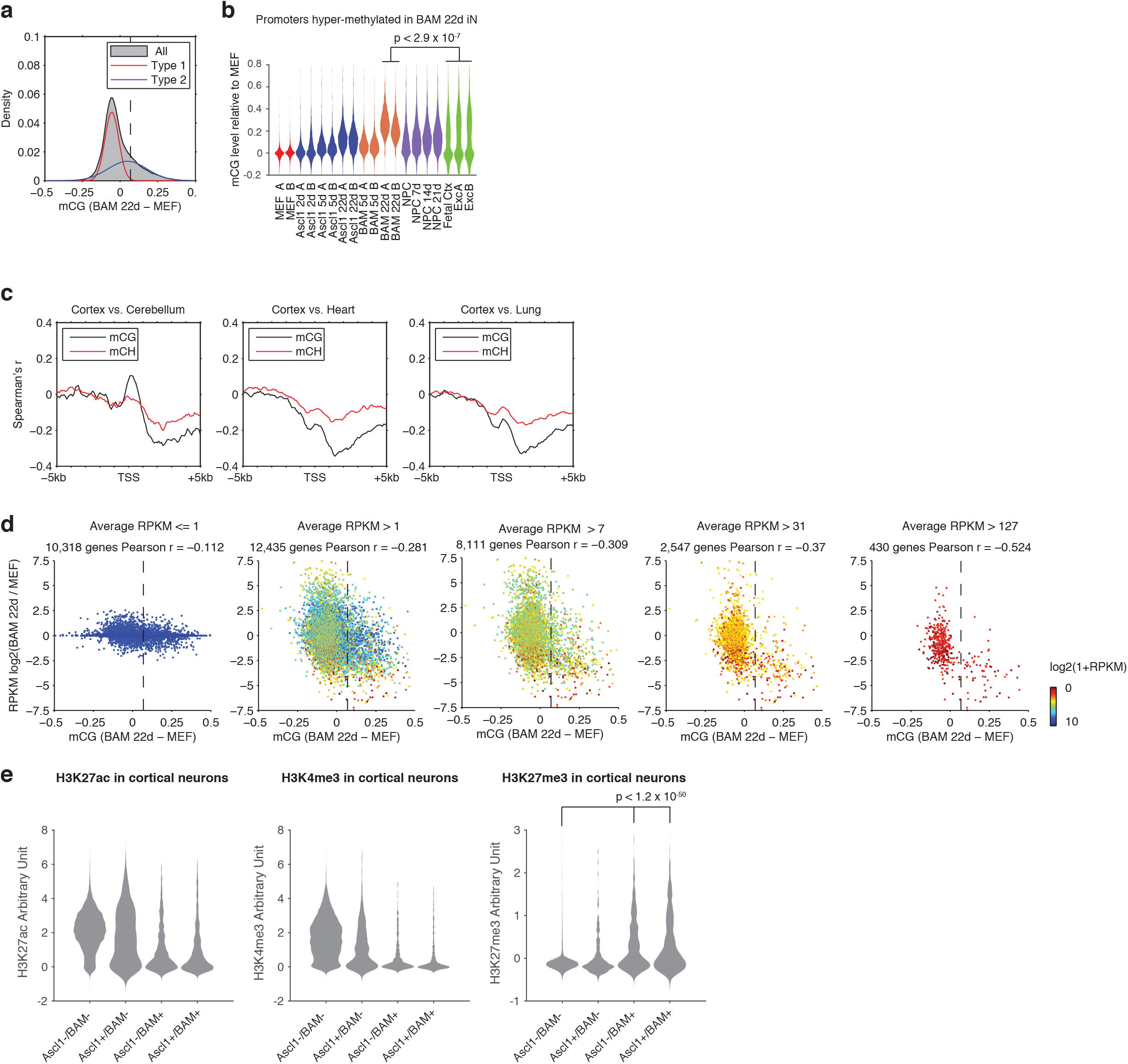
**a**, Probability density of normalized mCG changes (BAM 22d – MEF) for 1kb regions surrounding TSS. A Gaussian mixed model with two components was fitted to the data and resulted in two distributions (type A and B) with μ values of −0.0551 and 0.0485, σ values of 0.0021 and 0.0169 and weights of 0.5559 and 0.4441, respectively. The threshold of hyper-mCG TSS was determined to be 0.0691. **b.** mCG level of promoters showing hyper-methylation during direct reprogramming induced by BAM, in *in vitro* and *in vivo* grown cell. mCG levels were normalized to that of MEF. **c**, Correlation between mCG change and differential gene expression between cortex, cerebellum, heart and lung in +/− 5kb regions surrounding TSS. **d**. Scatter plots of promoter mCG change against differential gene expression between BAM 22d iN cells and MEF for genes with average RPKM <= 1, > 1, >7, > 31 and > 127. The vertical dashed line indicates the threshold of promoters showing significant hyper CG methylation. There is a strong inverse correlation between promoter hyper CG methylation and gene expression change. **e.** All promoters were separated to four categories depending on their hyper-mCG states in Ascl1 22d iN or BAM 22d iN cells. Violin plots show H3K27ac, H3K4me3 and H3K27me3 levels of the four types of promoters in cortical neurons to explore their chromatin states in *in vivo* grown cells.

**Supplementary Figure 4.**
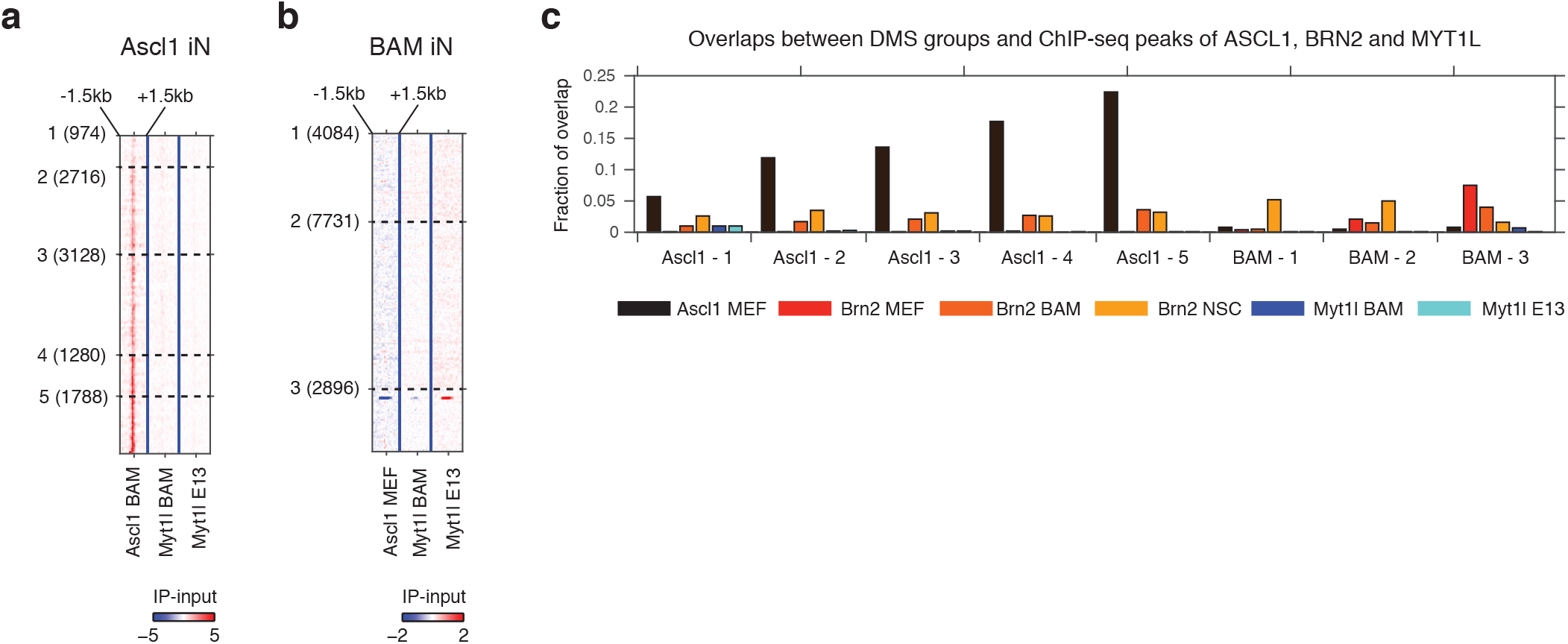
**a.** Signal intensity of Ascl1 and Myt1l ChIP-seq were plotted for 3 kb regions surrounding DMSs induced by Ascl1-only reprogramming. **b.** Signal intensity of Ascl1 and Myt1l ChIP-seq were plotted for 3 kb regions surrounding DMSs induced by BAM reprogramming. **c.** The overlap between CG-DMS groups and Ascl1, Brn2 and Myt1l binding sites.

**Supplementary Figure 5.**
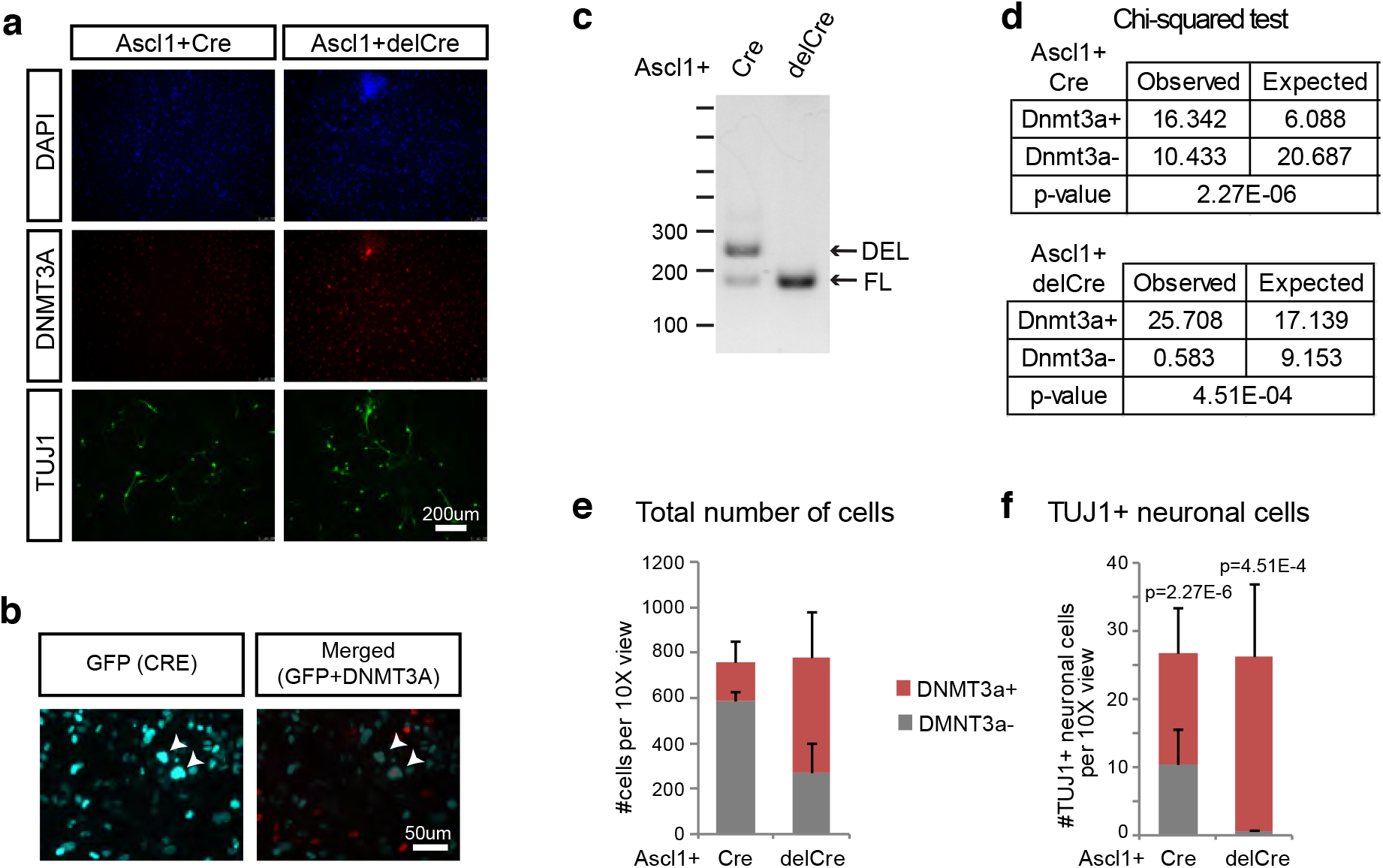
**a,** 10X view of immunostaining in Ascl1+CreGfp/delCreGfp samples. **b**, Corresponding GFP staining (for CREGFP fusion protein) for Fig. 5c. Even though iN cells are expressing high levels of CRE, which should loop out the Dnmt3a locus, cells are still expressing DNMT3A protein. **c,** Genotyping of Dnmt3a/b fl/fl cells. Floxed mutant bands (FL) appear at ~200bp while deleted bands (DEL) appear at ~250bp. **d**, Chi-squared test for significance for Suppl. Fig. 5e-f. The null hypothesis is that if Dnmt3a- cells reprogram as well as the Dnmt3a+ cells, there should be no significant difference in the frequency of Dnmt3a- cells and the frequency of cells that actually reprogram into neurons. There is a significant difference for both Cre (p=2.27E-6) and delCre (p=4.51E-4) conditions, indicating that Dnmt3a- cells are less efficient in reprogramming into iN cells. **e-f**, Average counts of DAPI+ cells **(e)** or TUJ1+ iN cells **(f)** 14d post-Ascl1 induction that are either DNMT3A positive (red) or negative (gray) per 10X field of view. Using a Chi-squared test (Suppl. Fig. 5d), there is a significant difference between the frequencies of total number DNMT3A- cells compared to the number of DNMT3A- iN cells in both Cre and delCre conditions.

